# The homeobox transcription factor Cux1 coordinates postnatal epithelial developmental timing but is dispensable for lung organogenesis and regeneration

**DOI:** 10.1101/2024.03.26.586673

**Authors:** Barbara Zhao, Jacob Socha, Andrea Toth, Sharlene Fernandes, Helen Warheit-Niemi, Brandy Ruff, Gurgit K. Khurana Hershey, Kelli L. VanDussen, Daniel Swarr, William J. Zacharias

## Abstract

Lung epithelial progenitors use a complex network of known and predicted transcriptional regulators to influence early lung development. Here, we evaluate the function of one predicted regulator, Cux1, that we identified from transcriptional regulatory analysis of the SOX9^+^ distal lung progenitor network. We generated a new Cux1-floxed mouse model and created an epithelial-specific knockout of Cux1 using Shh-Cre (Cux1^ShhCre-LOF^). Postnatal Cux1^ShhCre-LOF^ animals recapitulate key skin phenotypic features found in prior constitutive Cux1 knockout animals, confirming functionality of our new floxed model. Postnatal Cux1^ShhCre-LOF^ mice displayed subtle alveolar simplification and a transient delay in alveologenesis without persistent lung phenotypes or alterations in lung epithelial cell allocation. Cux1^ShhCre-LOF^ mice developed failure to thrive in their second and third weeks of life due to delayed ileal maturation, which similarly resolves by postnatal day 35. Finally, we challenged Cux1^ShhCre-LOF^ with influenza-mediated lung injury to demonstrate that Cux1^ShhCre-LOF^ mice undergo productive alveolar regeneration that is indistinguishable from WT animals. Together, these findings indicate that epithelial-specific loss of Cux1 leads to transient developmental delays in the skin, lung, and intestine without defects in definitive organogenesis. We conclude that Cux1 function is required for temporal optimization of developmental maturation in multiple organs with implications for susceptibility windows in developmental disease pathogenesis.

**One-Sentence Summary:** Deletion of key DNA binding domains leads to loss of Cux1 function in the lung, intestine, and skin characterized by transient failure to thrive without significant adult disease.

## Introduction

Lung development proceeds through a stereotypical series of morphogenetic processes to pattern both the proximal and distal lung compartments. During lung development, lung progenitor cells that express SOX9 and ID2 participate in early airway patterning and become restricted progenitors for the alveolar epithelium, extending distal lung saccules and generating both alveolar type 1 (AT1) and alveolar type 2 (AT2) cells. This progenitor activity is controlled by a combination of transcriptional regulators, including Sox9^2,3^, Id2^4^, Etv5^5^, Nkx2-1^6,7^, Foxp1/2^8^, and Foxa1/2^9^, which coordinate to balance proliferation and differentiation of this cell population. Recently, we reported a core transcriptional regulatory network (TRN) for this cell state, generated from single-cell (sc)RNA and scATACseq data used to train the paired expression and chromatin accessibility (PECA) model^10,11^. This TRN contained known regulators of SOX9^+^ distal lung progenitor function, as well as unknown potential regulators. We have validated one prediction from this regulatory model, which implicated phosphoinositide-3-kinase (PI3K) signaling in distal lung specification, but several other predicted regulators have not yet been carefully studied to date^11^. We posited that this TRN contains other important regulators of SOX9^+^ distal lung progenitor function. One attractive target was Cux1, a homeodomain transcription factor (TF)^12^ predicted to function as a core regulator in the SOX9^+^ distal lung progenitor network and previously implicated as a regulator of embryonic pulmonary development.

The Cux1 genomic locus is comprised of Cut repeat 1 (CR1), Cut repeat 2 (CR2), Cut repeat 3 (CR3), and the Cut homeodomain (CHD), all of which may function as DNA-binding domains^13,14^, in combination with two repressive domains at the C-terminus^15^. Post-translationally, Cux1 has been reported in up to six protein isoforms, denoted by their molecular weights as p200, p150, p110, p90, p80, and p75^16,17^. The majority of these protein isoforms are generated via cathepsin L-induced proteolytic processing of the full-length p200 isoform^18-20^; the biological function of some isoforms has been challenged^20,21^. The predominant isoforms of Cux1 (p110, p90, and p80) are known to function primarily as transcriptional repressors via either direct competition for binding site occupancy or recruitment of histone deacetylases via the C-terminal repressive domains^15,22,23^.

Cux1 activity is implicated in cell-cycle regulation^24-27^, cell migration^28-32^, and epithelial-mesenchymal transition (EMT)^33,34^ in a variety of tissues, among which include the kidney, pancreas, and lung. Interestingly, Cux1 also influences cellular senescence^35,36^ and base excision repair^37,38^. Cux1 expression is mediated by multiple signaling pathways, such as TGFβ^28,30,39-41^, Wnt^31,42,43^, and NF-_κ_B^44,45^, which effect its cellular functions in collagen synthesis, fibrosis, cell motility, EMT, and macrophage specification. In multiple cancer types, Cux1 paradoxically acts as both an oncogene and tumor suppressor; the current paradigm proposes that one Cux1 allele is inactivated to promote tumor initiation and the second allele is subsequently amplified to promote tumor progression^37,43,46-52^. Cux1 thus regulates a myriad of cellular functions to induce cell fate changes and tissue reorganization.

To date, three Cux1 germline knockout (KO) mouse models have been generated to study Cux1 function. The first model inserted a STOP codon into the CR1 exon, resulting in alternative splicing that generated normal amounts of the Cux1 protein all lacking CR1. These animals displayed wavy hair, curly whiskers, and impaired lactation without apparent pulmonary or endodermal phenotypes^53^.

A second model inserted a STOP codon into the CHD, resulting in a Cux1 protein lacking the CHD and C-terminal regions. These animals displayed failure to thrive, wavy whiskers, missing fur, and increased bacterial infections. The model was analyzed independently by two groups, with one group finding that homozygous KO mice were born at Mendelian ratios but died within a week^54^ and the other suggesting Cux1-null mice were born at non-Mendelian ratios^55^; these differences have been attributed to differential background strains. The third Cux1 constitutive knockout model replaced exons 22 and 23 with an in-frame lacZ gene, creating a functionally null fusion protein comprised solely of the N-terminal region through the start of CR3. The majority of inbred homozygous mice from this KO model died of “respiratory failure” shortly after birth with suggested delay in lung organogenesis and thickened alveolar septa lacking mature AT1 and AT2 cells^56^. Homozygous outbred KO mice survived past birth, but their lungs displayed alveolar simplification and decreased alveolar complexity compared to those of their WT littermates. Consistent with previous KO models, these homozygous mice presented with disrupted follicle morphogenesis and a complex skin phenotype^56^.

Despite these extensive prior reports, both the heterogeneity of these previous models and the lack of cell specificity limit our understanding of the role of Cux1 in lung biology. These published data and our TRN inference model provided strong rationale for our hypothesis that Cux1 function is critical for lung epithelial development. Therefore, we generated a novel Cux1-floxed mouse model to evaluate lung epithelial requirements for Cux1. Here, we show that, surprisingly and in contrast to prior findings, epithelial Cux1 expression is dispensable for lung development and regeneration following viral infection and is instead exclusively associated with transient developmental delays and failure to thrive.

## Results

### Rationale for Cux1 function in respiratory endoderm

We evaluated previously published data^11^ yielded from the paired expression and chromatin accessibility (PECA) model, which infers TRNs across diverse cellular contexts based (1) localization and activity of chromatin regulators with cis-regulatory elements, (2) sequence-specific transcription factor binding site prediction, and (3) gene expression of putative targets at the RNA level^10^. The PECA model identified CUX1 as one of the top 3 predicted regulators of gene expression in SOX9^+^ distal lung progenitors, ranked immediately after known key regulators ETV5 and ID2 (Figure 1A), with predicted regulated gene targets highly enriched in lung epithelial development-associated GO Biological Process terms. Interrogation of LungMAP scRNAseq data^57^ demonstrated that Cux1 expression is enriched in epithelial cells, especially AT1 cells, across the murine lifespan (Figure 1C) and peaks during late embryonic development. Immunohistochemistry (IHC) showed CUX1 expression in the distal lung at E16.5, with protein localizing primarily to the developing distal lung saccular epithelium (Figure S1A-C).

**Figure 1.**
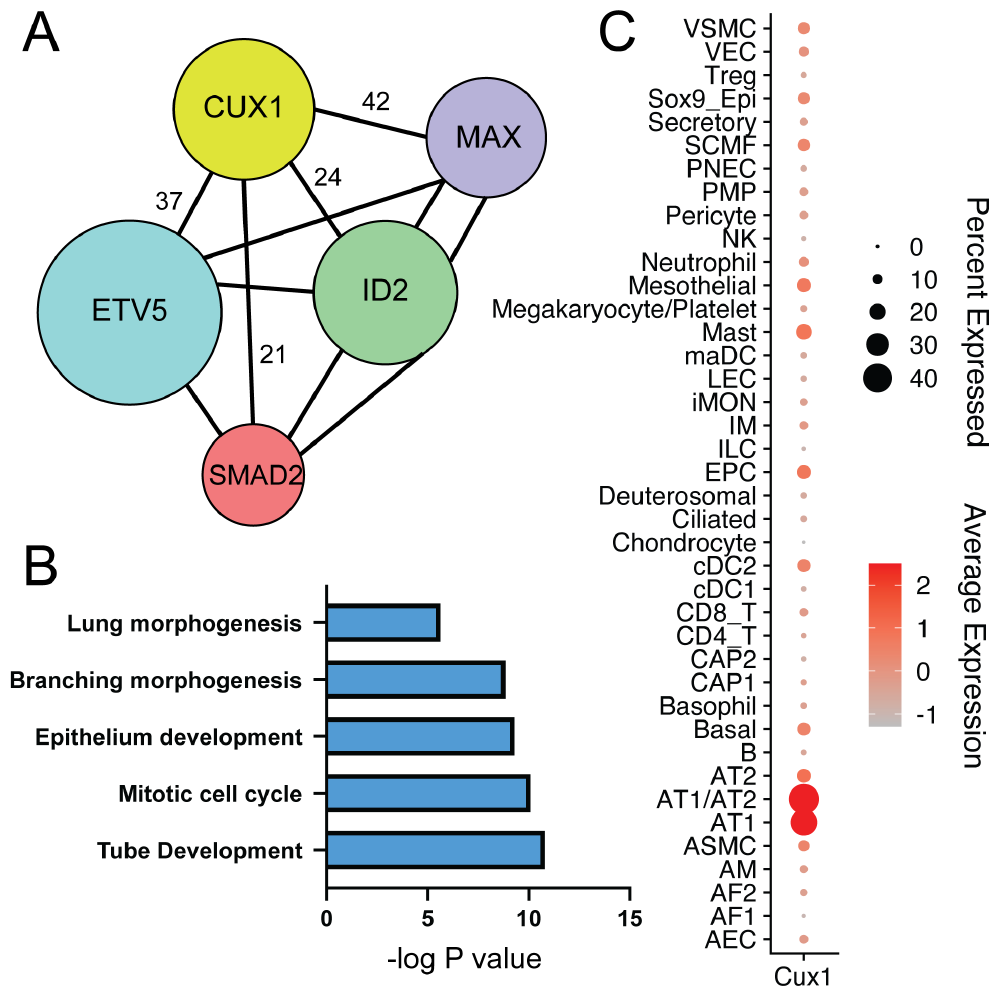
Evidence suggests Cux1 functions in lung epithelial development. (A) Transcriptional regulatory network (TRN) showing interaction of top 5 TFs in PECA analysis for E16.5 SOX9^+^ distal lung progenitors. Size of circle represents the number of targets in predicted SOX9^+^ network, and numbers next to spokes emanating from CUX1 denote the number of co-regulated targets predicted for each of the other TFs. (B) GO Biological Process term enrichment of predicted CUX1 gene targets in SOX9^+^ cells as determined by ToppGene^1^. Enrichments are presented as -log of the corrected p-value. (C) Cux1 gene expression scRNA seq data from the LungMAP consortium CellRef dataset spanning E16-PND28. Cux1 is highly expressed in AT1 and AT1/AT2 cells across the murine lifespan, with absolute highest expression at embryonic timepoints.

### Generation of a conditional Cux1 loss of function mouse model

Learning from prior experience with Cux1 germline KO models, we created a new mouse model by generating genomic knock-in of two loxP sites flanking exons 22 and 23, which contain the critical DNA-binding CR3 and CHD domains of the native Cux1 locus (Figure 2A). Recombination of this allele removes exons 22 and 23, closely recapitulating genomic replacement of these same domains with LacZ in the most severe of the constitutive KO models^56^ and impacting all known protein isoforms of Cux1 (Figure 2B). In addition, a second targeting event was used to knock-in an in-frame HA tag at the 3’ end of exon 24, adding the tag to the C-terminus of the protein for detection of all known isoforms of Cux1 produced by differential splicing or enzymatic cleavage via cathepsin L. Sanger sequencing verified successful targeting of the Cux1 locus for both loxP sites and the terminal HA tag (data not shown).

**Figure 2.**
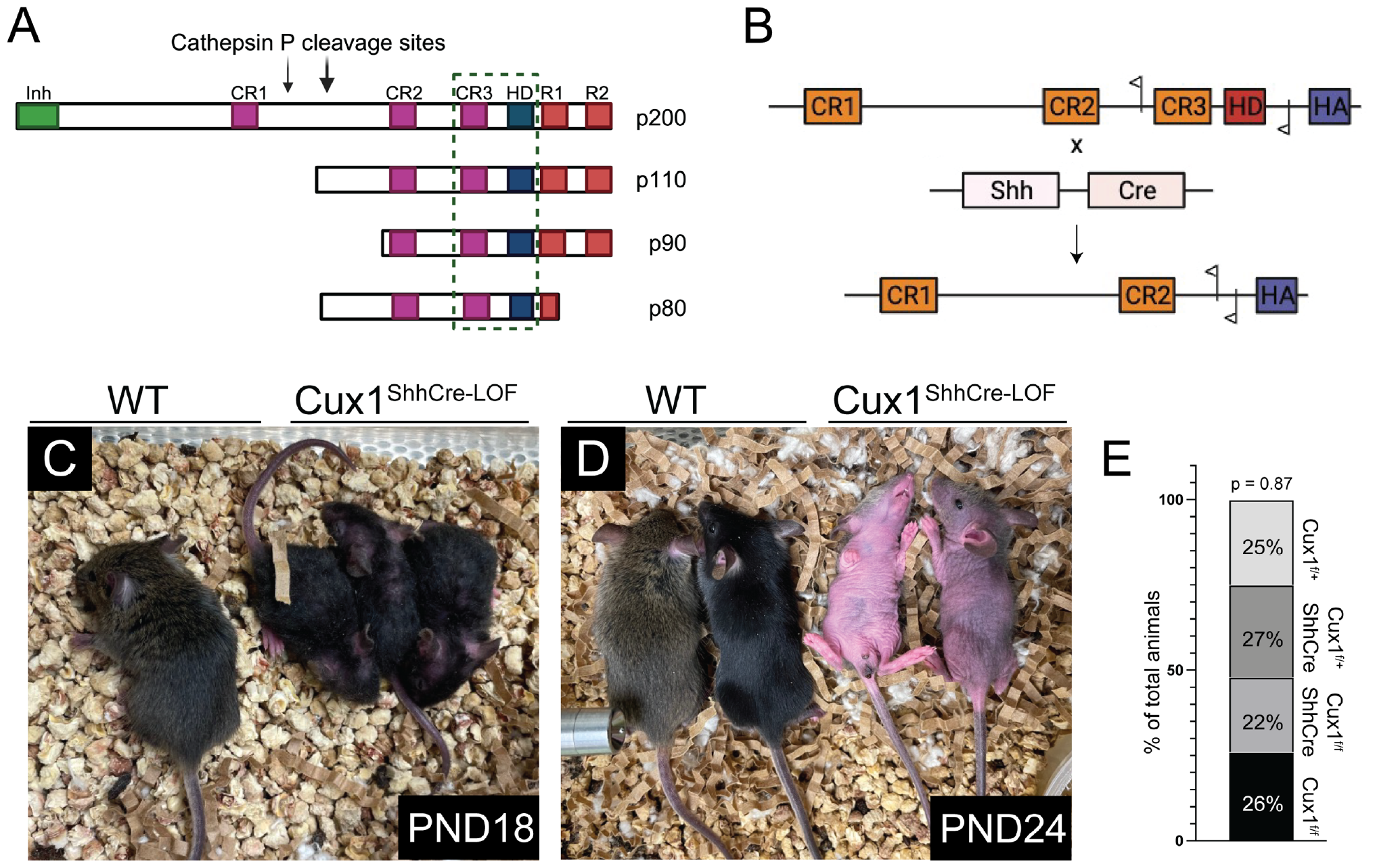
Construction and gross phenotype of Cux1 conditional deletion mouse line. (A) Schematic of major Cux1 protein isoforms, highlighting the consistent presence of Cut repeat 3 (CR3) and the homeodomain (HD) in all major isoforms. Dotted box shows area of protein excised following loss of exons 22 and 23 by Cre-Lox recombination. (B) Schematic of 3’ end of Cux1 locus in knockout animals, showing floxed allele before (top) and after (bottom) Cre-mediated recombination via Shh-Cre. (C-D) Gross phenotype of Cux1^ShhCre-LOF^ animals at PND18 (C) and PND24 (D), highlighting smaller size and range of skin abnormalities, including patchy coats (C) and significant loss of hair (D). (E) Percentage of animals of each phenotype observed in breeding of Cux1^ShhCre-LOF^ animals, with no significant difference between groups by two-sided Fisher’s Exact Test from N = 172 animals across 8 litters. *(Inh = inhibitory domain; CR = Cut repeat domain; HD = homeodomain, R = repressor domain; HA = HA tag; WT = wild type)*.

To generate Cux1 loss of function (LOF) in the developing lung epithelium, Cux1^fl/fl^ animals were crossed to *Shh*^*Cre*^ mice to produce Cux1^ShhCre-LOF^ mice (Figure 2B). Shh-Cre is expressed in the developing endoderm and ectoderm^58,59^ and is commonly used to generate epithelial-specific knockouts in pulmonary biology. Unexpectedly, Cux1^ShhCre-LOF^ mice were born at Mendelian ratios and survived transition to air breathing (Figure 2E). During postnatal development, Cux1^ShhCre-LOF^ developed failure to thrive, with reduced weight gain (Figure 3) and grossly apparent skin abnormalities (Figure 4). By postnatal day (PND) 18, Cux1^ShhCre-LOF^ demonstrated wavy pelage and loss of fur compared to their WT littermates (Figure 2C-D); these findings are similar to reported skin abnormalities in constitutive Cux1 KO models.

**Figure 3.**
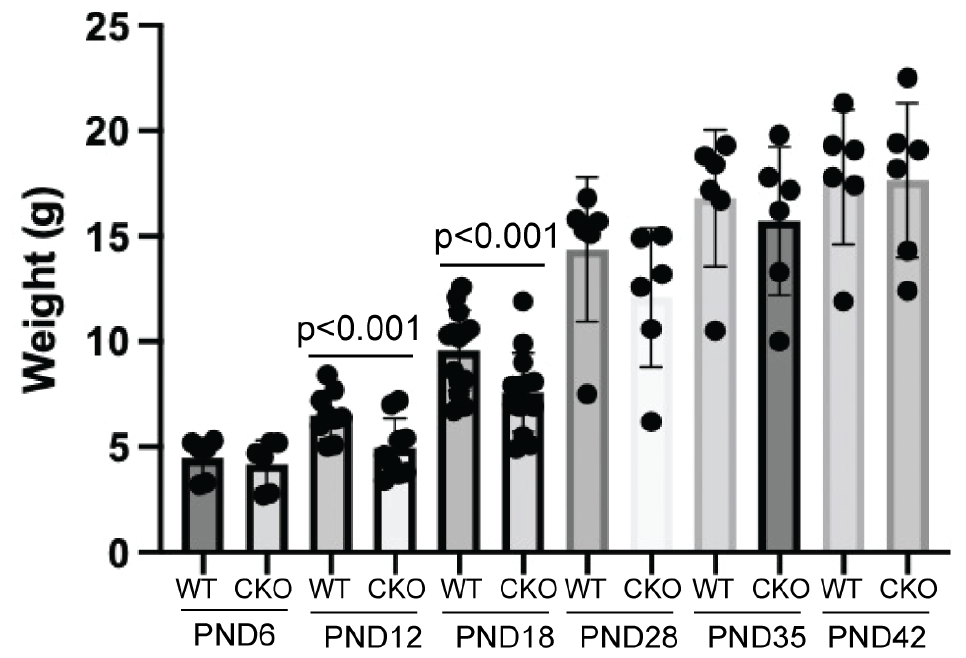
Cux1^ShhCre-LOF^ animals demonstrate failure to thrive, which resolves by the 4^th^ week of life. Cux1^ShhCre-LOF^ animals exhibit no differences in weight gain at PND6 compared to their WT littermates, followed by a delayed weight gain across their 2^nd^ and 3^rd^ weeks of life.

**Figure 4.**
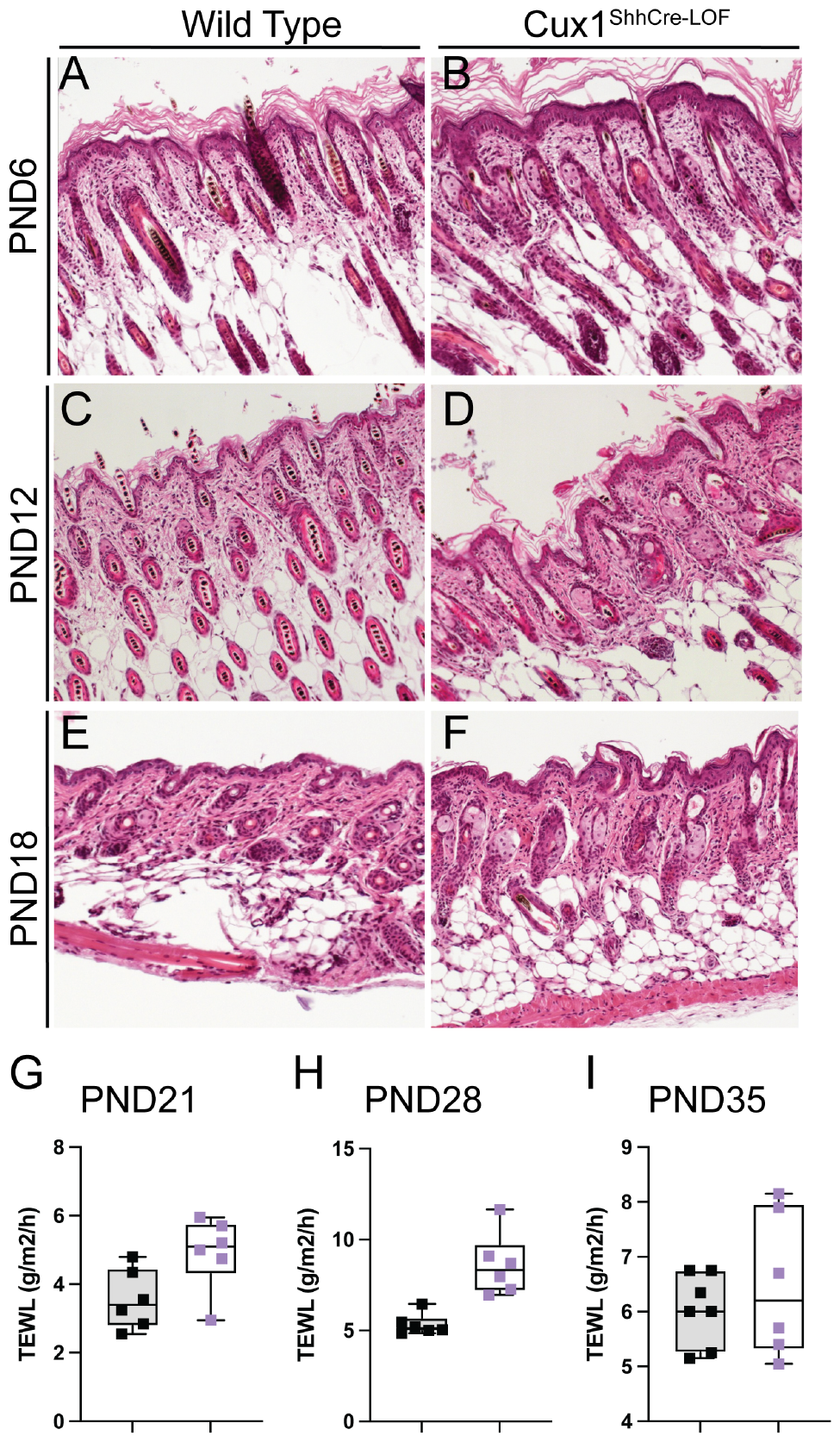
Postnatal Cux1^ShhCre-LOF^ mice have disorganized skin development and impaired barrier function. (A-F) Cux1^ShhCre-LOF^ mice demonstrate progressively disorganized, sclerotic hair follicles at PND12 and PND18 compared to their WT littermates. (G, H) Trans-epidermal water loss (TEWL) measurements taken at PND21 and PND28 show that Cux1^ShhCre-LOF^ mice have significantly impaired skin barrier function compared to their WT littermates. (I) TEWL measurements for Cux1^ShhCre-LOF^ mice and their WT littermates at PND35 reveal no significant differences, suggesting that skin barrier function impairment of Cux1^ShhCre-LOF^ animals is transient. Black = WT, purple = Cux1^ShhCre-LOF^.

### Cux1^Shh-Cre^ mice present with disorganized hair follicles and impaired skin barrier function

Next, we more carefully evaluated the skin phenotype in Cux1^ShhCre-LOF^ animals. Previous Cux1 KO mouse models detailed a skin phenotype consisting of wavy pelage, loss of pelage, and aberrant hair follicle development. Histological evaluation of the skin Cux1^ShhCre-LOF^ mice at PND6, PND12, and PND18 demonstrated disorganized and sclerotic hair follicles compared to their WT littermates (Figure 4A-F). We then assessed skin barrier function in Cux1^ShhCre-LOF^ mice and their WT littermates via measurement of trans-epidermal water loss (TEWL), which quantifies the amount of water lost across the stratum corneum due to passive evaporation^60^. Increased TEWL measurements correspond with decreased skin barrier function. We found that skin barrier function was significantly impaired in Cux1^ShhCre-LOF^ mice compared to their WT counterparts at PND21 and PND28, and that skin barrier impairment resolved by PND35 (Figure 4G-I). We noted contemporaneous regrowth of pelage in Cux1^ShhCre-LOF^ after PND28. These findings suggest that Cux1^ShhCre-LOF^ mice present with significant developmental abnormalities in epidermal growth, leading to impaired epidermal barrier function that at least partially resolves during the 5^th^ week of life. These findings recapitulated the key skin phenotypes of previous KO animals, confirming that the Cux1^fl/fl^ allele recombined as expected to generate a functionally null CUX1 allele.

### Postnatal Cux1^ShhCre-LOF^ mice develop subtle alveolar simplification during alveologenesis

We next turned our attention to evaluating the respiratory development of Cux1^ShhCre-LOF^ animals. To determine the impact of epithelial deletion of Cux1 in respiratory development, we generated a developmental time series, which began at the saccular stage (E18.5) and continued throughout alveologenesis (PND6, PND12, and PND18). (Figure 5). At E18.5, Cux1^ShhCre-LOF^ animals displayed no apparent histological phenotype (Figure 5A), suggesting that epithelial Cux1 expression was not functionally required for embryonic lung development. However, at PND6, Cux1^ShhCre-LOF^ mice displayed subtle alveolar simplification compared to their WT littermates (Figure 5B,F), with differences in morphological measurements of mean linear intercept (L_m_), volume density of alveolar septa (V_vsep_), mean trans-sectional alveolar wall length (L_mw_), and surface area density of airspaces (Sv_air_) (Figure 5I-L). These differences improved at PND12 and resolved by PND18, suggesting only a brief delay in alveologenesis in Cux1^ShhCre-LOF^ animals.

**Figure 5.**
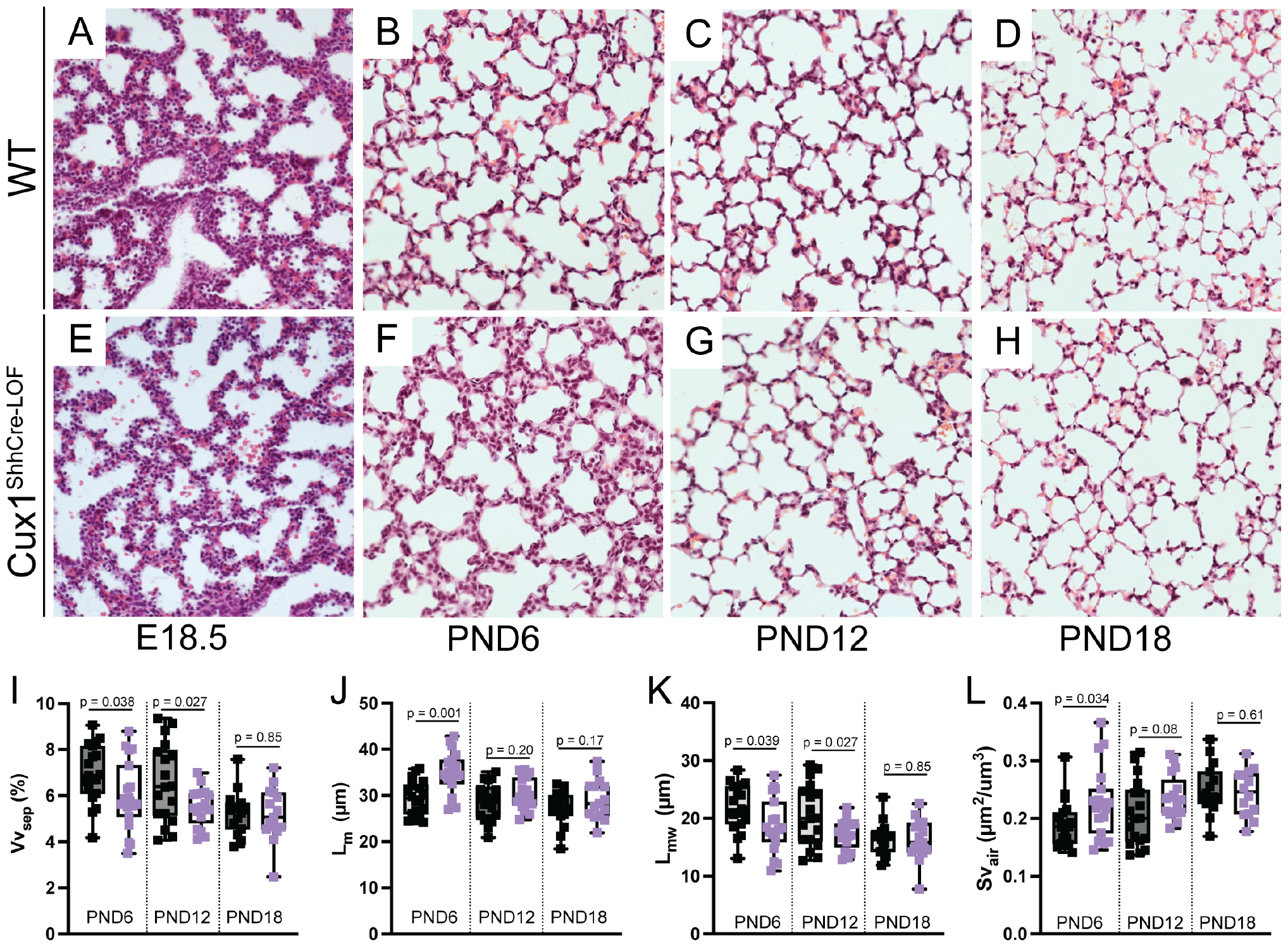
Postnatal Cux1^ShhCre-LOF^ mice develop subtle alveolar simplification during alveologenesis. (A-H) Cux1^ShhCre-LOF^ mice demonstrate alveolar simplification that is grossly notable at PND6 (B,F) and resolves by PND18 (D,H). (I-L) Quantification of lung morphometry with volume density of alveolar septa (V_vsep,_ I), mean linear intercept (L_m_, J), mean trans-sectional alveolar wall length (L_mw_, K), and surface area density of airspaces (Sv_air_, L) demonstrates significant differences in all measures at PND6, slight differences at PND12, and no difference at PND18. P values calculated with two-tailed student’s t tests comparing Cux1^ShhCre-LOF^ to WT littermates at the same time point. Black = WT, purple = Cux1^ShhCre-LOF^. Images taken with 20x objective.

We next evaluated whether absence of Cux1 function led to differences in the cell fate acquisition of lung epithelial cells during alveologenesis. IHC confirmed loss of HA-tagged protein expression in Cux1^ShhCre-LOF^ lungs in both AT1 and AT2 cells during alveologenesis at PND6. No differences in AT2 cell number or AT1 cell surface were evident despite clear loss of CUX1 expression and alveolar simplification, suggesting that CUX1 is not required for lung epithelial fate specification (Figure 6A-D). Alveolar simplification can result from failure to thrive, suggesting an additional mechanism for the delayed respiratory development seen in Cux1^ShhCre-LOF^ mice. We therefore turned our attention to characterizing the intestinal development of Cux1^ShhCre-LOF^ animals

**Figure 6.**
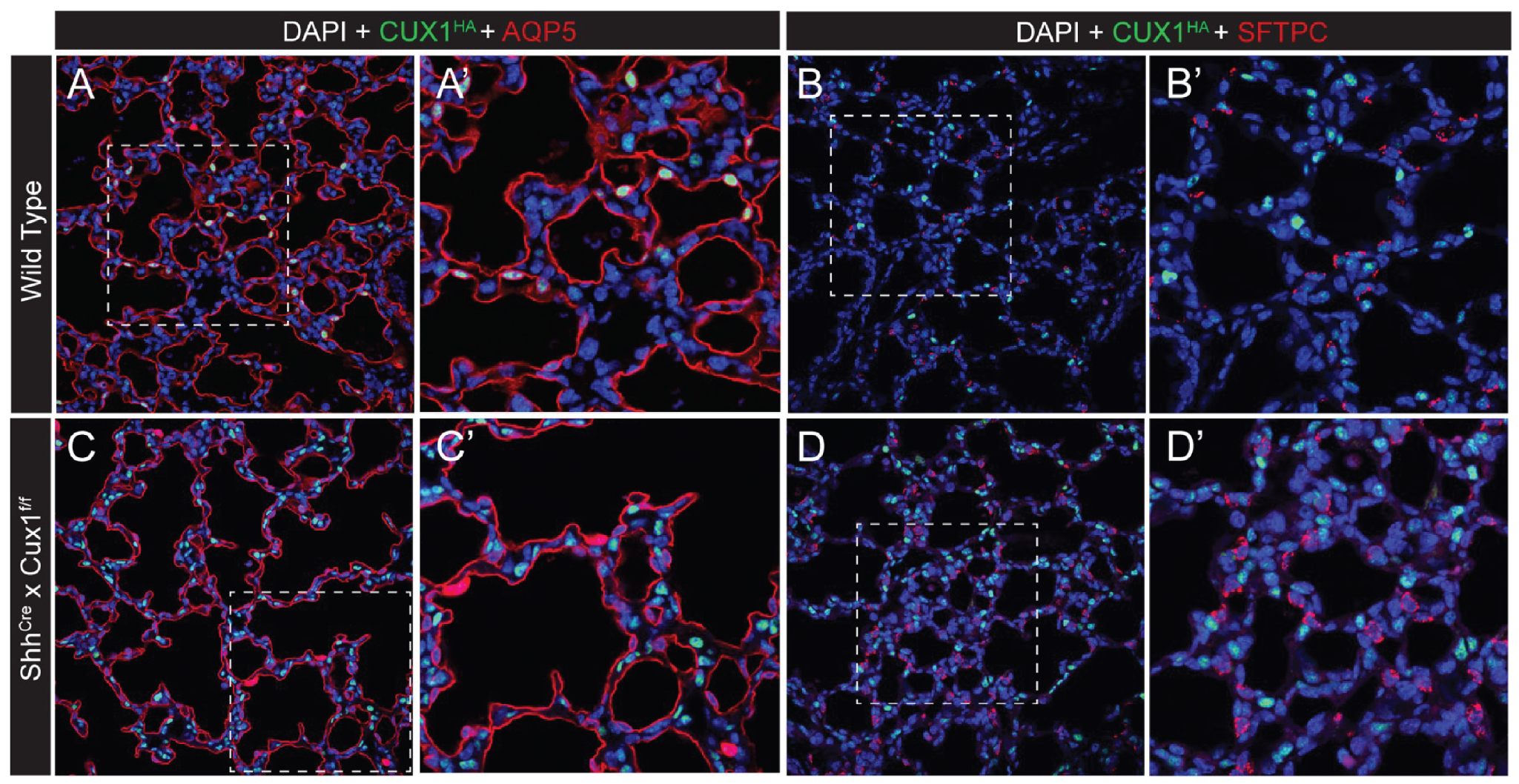
Cux1^ShhCre-LOF^ mice appropriately develop mature alveolar type 1 and alveolar type 2 cells. (A-B) WT mice demonstrate expression of HA-tagged CUX1 in the nuclei of both AT1 (A) and AT2 (B) pneumocytes at PND6. (C-D) CUX1-HA expression is lost in both AT1 and AT2 cells, but no significant change in either cell population is notable despite subtle alveolar simplification. All images taken at 20x and 60x. *(AQP5 = Aquaporin 5, AT1 cell marker; SFTPC = Surfactant Protein C, AT2 cell marker*.

### Postnatal Cux1^ShhCre-LOF^ animals fail to thrive and present with developmental delays in the ileum

As noted above, Cux1^ShhCre-LOF^ mice were visibly smaller than their WT counterparts (Figure 2D-E, Figure 3). Cux1^ShhCre-LOF^ mice weigh significantly less than WT littermates at PND12 and PND18 (Figure 3), consistent with the failure to thrive phenotype. As developmental delays in gastrointestinal (GI) development are principal causes of failure to thrive and Shh-Cre is expressed throughout the intestinal epithelium^61,62^, we hypothesized that Cux1^ShhCre-LOF^ mice might have altered intestinal development.

We therefore evaluated the duodenum, jejunum, ileum, colon, and stomach of Cux1^ShhCre-LOF^ mice and their WT littermates. We confirmed that the HA-tagged Cux1 protein was absent from the gut epithelium of Cux1^ShhCre-LOF^ mice and broadly expressed in both the gut mesenchyme and epithelium of WT littermates at PND18 (Figure S2D,I). Histological samples of the duodenum, jejunum, and colon demonstrated no distinguishable differences between WT and Cux1^ShhCre-LOF^ mice (Figure S2A-J). Stomach glands of Cux1^ShhCre-LOF^ mice were slightly shorter than those of their WT littermates (Figure S2A,F). The major phenotypic change notable in the intestines of Cux1^ShhCre-LOF^ mice was a significant increase in ileal epithelial vacuoles at PND18 (Figure 7A-G, S2D,I), with no significant differences in mean ileal crypt depth or villus height (Figure 7H-I). Concordantly, we observed no differences in ileal crypt proliferation (Figure S3G-H).

**Figure 7.**
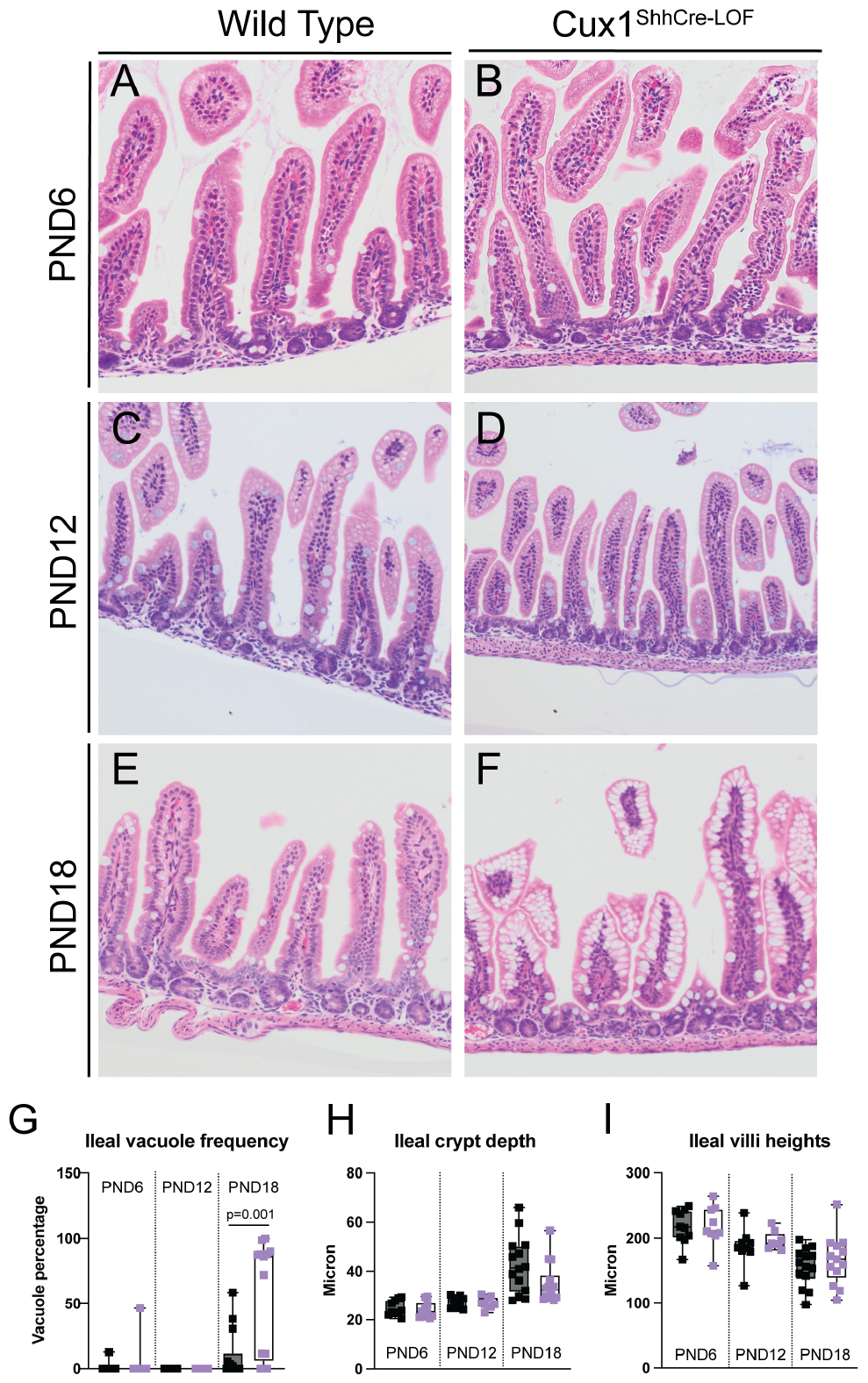
Postnatal Cux1^ShhCre-LOF^ mice exhibit developmentally delayed gut maturation. (A-F) The ileum of Cux1^ShhCre-LOF^ mice is indistinguishable from the ileum of their WT littermates at PND6 (A-B) and PND12 (C-D). By PND18, Cux1^ShhCre-LOF^ ileum display vacuoles that are not present in WT littermates. (G-I) Quantification of the frequency of villi with vacuoles (G), Ileal crypt heights (H), and ileal villi heights (I) in WT vs. Cux1^ShhCre-LOF^ mice. Villi and crypt counts made per 100 villi or 100 crypts.

We hypothesized that the ileal vacuoles observed in Cux1^ShhCre-LOF^ mice may contain lysosomal enzymes as seen in early ileal development^63,64^ or mucous as observed in foveolar gastric metaplasia^65^. However, Alcian blue staining for mucin (Figure S3A-C) and Oil Red O staining for lipids (Figure S3D-E) reveal that these ileal vacuoles contain neither mucous nor lipids, which made these possibilities unlikely. An alternative interpretation arose from prior literature. During the suckling stage of gut maturation, pinocytotic vesicles transport dietary proteins and immunoglobulins through the stomach and intestinal epithelium^66^. While epithelial cells transition to mature intestinal enterocytes, these pinocytotic vesicles are degraded by high levels of lysosomal enzymes within the developing villus epithelium^67^. We therefore conclude that the ileal epithelium in Cux1^ShhCre-LOF^ is immature at PND18, leading to the failure to thrive phenotype. This conclusion is supported by the observation that ileal maturation appears complete by PND35 (Figure S5), with reduced vacuolization and intestinal morphology indistinguishable from WT littermates. This corresponds to a significant weight gain in the fourth week of life; by PND28, Cux1^ShhCre-LOF^ murine weight is no longer significantly reduced, and by PND42, their weight is indistinguishable from WT littermates (Figure 3). We therefore conclude that epithelial Cux1 function is involved in the optimization and timing of developmental processes in the skin, lung, and gut but is not required for definitive organogenesis in any of these organs.

### Cux1 is not required for lung epithelial regeneration and repair

Many regulators that are dispensable to early development in murine models may be critical for adult homeostasis, regeneration, and disease. Cux1 has been reported to be haploinsufficient in many cancers, including lung cancer, and is associated with overexpression in tumors with high proliferative and malignant potential^46,47^. We therefore reasoned that Cux1 may be required for adult progenitor function in the lung. Cux1^ShhCre-LOF^ animals live to adulthood, so we administered adult a mild dose of intranasal H1N1 influenza (strain PR8) to Cux1^ShhCre-LOF^ animals as an infectious challenge to assess lung regenerative capacity. Weight trends in Cux1^ShhCre-LOF^ animals were indistinguishable from their WT littermates, and no animals in either group died as result infection (Figure S4A). We evaluated histology and lung injury score at 28 days post infection (DPI) and observed no discernable differences in the extent or histological phenotype of epithelial injury between Cux1^ShhCre-LOF^ and WT animals. (Figure 8A-B). Alveolar regeneration with appropriate differentiation of AT1 and AT2 cells was present in Cux1^ShhCre-LOF^ animals, and there was no difference in epithelial proliferation as marked by Ki67 expression (Figure 8C-D). We noted similar degrees of AT2 hyperplasia in both Cux1^ShhCre-LOF^ mice and their WT littermates. These findings suggest that Cux1^ShhCre-LOF^ mice retain major epithelial patterns of regeneration following influenza-induced injury. We therefore conclude that epithelial CUX1 is not required for lung regeneration or repair following acute viral injury.

**Figure 8.**
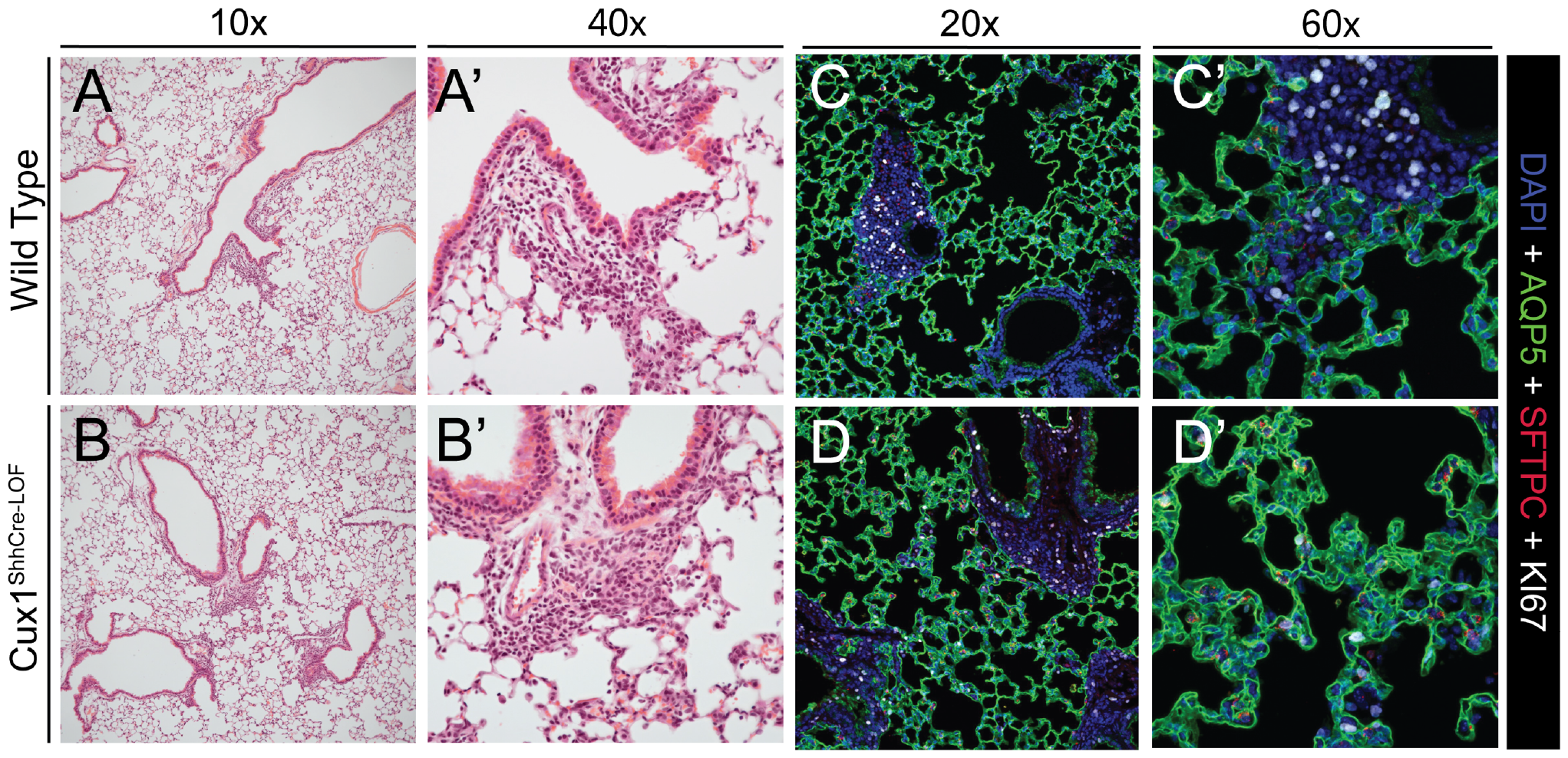
Cux1^ShhCre-LOF^ mice display intact lung injury resilience and functional regeneration following influenza A infection. (A-B) 6–8-week-old Cux1^ShhCre-LOF^ animals and WT littermates develop similar levels of airway and alveolar epithelial injury following infection with PR8 H1N1 influenza at 28 DPI. (C-D) Both AT1 and AT2 cells regenerate appropriately in Cux1^ShhCre-LOF^ animals at 28 DPI. No differences in epithelial proliferation are evident as measured by KI67 expression.

## Discussion

Transcriptional regulatory network inference is a powerful tool used to generate testable hypotheses about gene regulation from increasingly available single cell data. TRN inferences thus provide a pathway to study specific, complex mechanisms predicted by large scale explorative studies in lung biology. Recent work from our group and others has demonstrated that TRN inference can identify unexpected functions of even well-described transcriptional regulators^11,68^. Fundamentally, however, all TRN inferences generate predictions that require functional validation. Here, we show that despite extensive evidence implicating Cux1 as a key regulator of distal lung development, Cux1 function is largely dispensable for lung epithelial biology throughout development and regeneration following acute viral injury.

What does this surprising result imply about the validity of the original prediction from the TRN? Cux1 is likely involved in regulating lung epithelial development, as evidenced by the clear developmental delay seen during alveologenesis. Cux1 also participates in intestinal and skin development, as suggested by previous global knockout models and confirmed in present data from our new floxed model. However, these differences are partial, transient, and do not confer a longer-term susceptibility to epithelial injury or defect in repair. We therefore conclude that CUX1 likely functions as a cooperative TF functioning to maintain temporal dynamics^69^ in lung epithelial regulatory networks.

Transcriptional regulation is dynamic^70-72^, as multiple known TFs function primarily to stabilize or prolong the function of previously assembled gene expression complexes. Our TRN data implies that Cux1 may function in a similar way, given its extensive co-regulation of genes indispensable to distal lung epithelial cell development and coregulation of genes downstream of core TFs, such as Etv5 and Id2. Loss of Cux1 function is insufficient to disrupt the activity of these factors, as evidenced by the proper epithelial patterning observed in Cux1^ShhCre-LOF^ animals. However, the developmental delay observed in our study implies that Cux1 does promote the function of these core TFs, which may be critical for resiliency during development or in other contexts not yet evaluated in our study. Our hope is that the Cux1^fl/fl^ mouse will promote future studies to identify the core processes for which Cux1 is more central.

Finally, recent data highlights the concept of “susceptibility windows” in disease development, especially in the pediatric period^73,74^. Temporal control of gene expression may control the duration of such susceptibility windows^73^. Susceptibility windows during postnatal development have been recognized for lung disease^75^, atopic dermatitis^76,77^, and development of metabolic syndrome^78^, obesity, and functional and inflammatory bowel disease^79,80^. Given the contemporaneous delay in development of lung, skin, and gut in our study, it is tempting to speculate that factors such as CUX1 may optimize developmental timing and function to limit these critical windows, thereby providing resilience during postnatal development to environmental stimuli.

## Acknowledgements

The authors would like to thank the Bioimaging and Analysis Facility (especially director Matt Kofron) of the Cincinnati Children’s Research Foundation for technical support, as well as the CCHMC Pathology Research Core for tissue processing and embedding.

## Author Contributions

Conceptualization – BZ, AT, DS, WJZ

Data acquisition – BZ, JS, SF, HWN, BR

Data analysis – BZ, WJZ

Supervision – GKKH, KVD, DS, WJZ

Writing – original draft – BZ,

WJZ Writing – review and editing - all authors

## Data and materials availability

All data is available by request to the corresponding author.

## Materials and Methods

### Ethical compliance and animals

All animal studies were conducted under the guidance and supervision of the Cincinnati Children’s Hospital Medical Center (CCHMC) Institutional Animal care and Use Committee (IACUC) in accordance with CCHMC regulatory and biosafety protocols. Mouse lines used included: Cux1^fl/fl^, and Shh^Cre^. All experiments included both male and female mice. Controls in all cases included littermates lacking ShhCre expression.

### Mouse design

To create the Cux1^fl/fl^ mouse, we first knocked-in two loxP sites around CR3 and CHD, which are critical DNA-binding domains of Cux1, and added an HA tag in exon24 of the Cux1 gene for future molecular studies. The two loxP sites were knocked-in via Hygromycin and Neomycin resistance gene cassettes into murine embryonic stem cells (ESCs). Post-selection, these hybrid ESCs were microinjected into CD1 blastocysts. Resulting chimeras were bred with flippase (FLP) C57BL/6 mice to knock out the resistance gene cassettes. C57BL/6 mice with both loxP sites were then bred with *Shh-Cre* mice to produce Cux1^fl/fl^; Shh-Cre mice (Cux1^ShhCre-LOF^).

### Sanger sequencing

Ear punches from Cux1^fl/fl^ mice were lysed with a mixture of Lysis Buffer and Proteinase K. PCR was run on these samples with LOX1 and SQ1 primers for the first loxP site, SQ2 and SQ3 primers for the second loxP site, and specially designed F1 and R1 primers for the HA tag (Supplementary Table 1). Following PCR, bands were separated on a 1.5% agarose gel, cut from the gel via UV light visualization, and extracted from the gel with the QIAquick Gel Extraction Kit (Qiagen). DNA samples were submitted to the CCHMC Genomics Sequencing Facility for sequencing results.

**Supplementary Table 1.**
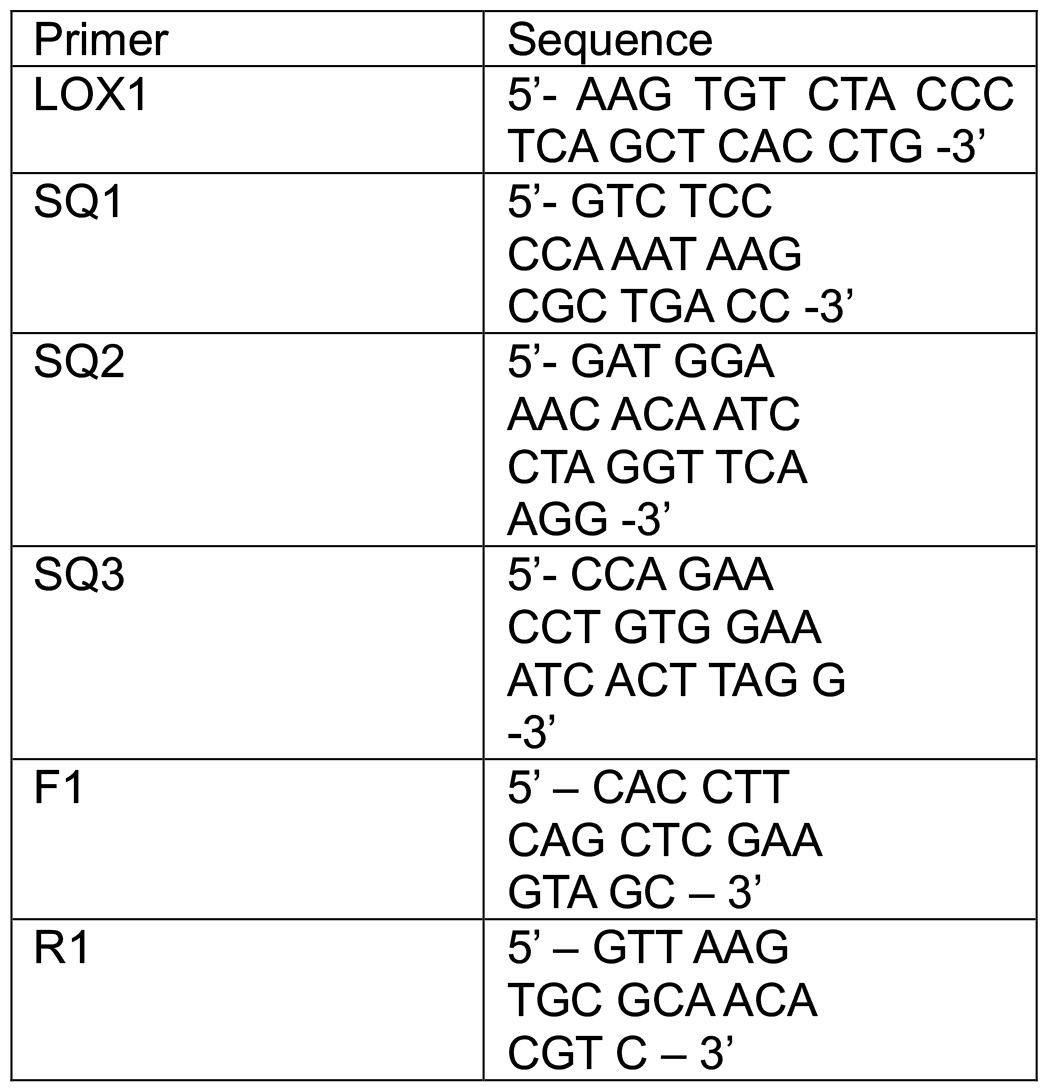
Primer sequences for loxP sites and HA tag sequencing.

### Mouse lung harvest

Mice harvested at E18.5 and postnatal days (PND) 6/12/18/35 were anesthetized via CO_2_ inhalation, followed by euthanasia via cervical dislocation and thoracotomy. Mice harvested in containment following PR8 H1N1 influenza infection were anesthetized via IP Ketamine + Xylazine, followed by euthanasia via cervical dislocation and thoracotomy. The chest cavity was opened to expose the heart and lungs. The right ventricle was perfused with 10mL of cold PBS to clear blood from the lungs. For tissue fixation for histology and immunofluorescence, the trachea was cannulated, and lungs were inflated via syringe using 4% paraformaldehyde (PFA). Inflated lungs were immersed in either a conical or small glass jar of 4% PFA and left on a rocker at 4°C overnight.

### Processing fixed lung tissue for histology and immunofluorescence

Following inflation, fixed lung tissue was trimmed and place into cassettes the next day. The cassettes were washed 3X in DEPC-treated PBS, 1X in DEPC-treated 30% ethanol, 1X in DEPC-treated 50% ethanol, and 1X in DEPC-treated 70% ethanol. Following a standardized overnight automated processing protocol (Thermo Scientific, Excelsior ES), the samples were embedded in paraffin. Samples were sectioned at a thickness of 5μm. Paraffin sections were incubated at 65 °C for 2 h, deparaffinized in xylene (3x for 10 min), rehydrated through an ethanol gradient, and standard H&E staining was performed. Slides were mounted with Permount Mounting Medium (Electron Microscopy Sciences, 17986-05) and cover-slipped with #1.5 Gold Seal 3419 Cover Glass (Electron Microscopy Sciences, 63790-01). Immunofluorescence on paraffin sections was performed as previously described^81^. Briefly following deparaffinization, rehydration, and sodium citrate antigen retrieval (10 mM, pH 6.0), and blocking, immunofluorescence was performed on paraffin sections using antibodies in Supplementary Table 2 and the following reagents: ImmPRESS® HRP Horse Anti-Rabbit IgG Polymer Detection Kit (Vector Labs, MP-7401-50). Following the application of TSA fluorophores (listed in Supplementary Table 2; 1:100), sections were stained with DAPI (Invitrogen, D1306; 1:1000) and mounted using Prolong Gold antifade mounting medium (Invitrogen, P36930).

**Supplementary Table 2.** Antibodies for immunofluorescence on paraffin sections.

| Primary antibody | Secondary detection |
| --- | --- |
| anti-HA (Rabbit, Cell Signaling Technologies – 1:800) | TSA Plus Cyanine 3.5 (Akoya Biosciences, NEL763001KT) |
| anti-Ki67 (Rabbit, Abcam, ab1667 – 1:200) | TSA Plus Cyanine 5 (Akoya Biosciences, NEL745001KT) |
| anti-Sftpc (Guinea pig, Seven Hills Bioreagents, GP992 – 1:500) | Goat anti-Guinea Pig IgG Alexa Fluor 555 (Invitrogen, A21435) |
| anti-Aqp5 (Rabbit, Abcam, ab92320 – 1:100) | TSA Plus Fluorescein (Akoya Biosciences, NEL741001KT) |
| anti-Cux1 (Rabbit, Abcam, ab73885 – 1:100) | TSA Plus Fluorescein (Akoya Biosciences, NEL741001KT) |

### Murine skin tissue harvest and fixation for histology and immunofluorescence

Concurrently with mouse lung harvest, skin samples were collected from PND6/12/18/35 mice. All skin samples were cut in a square from the center backside of each mouse following shaving of the area via razor blade. Skin samples were laid flat on a 3% agarose gel plate and pinned to the agarose in all 4 corners. Skin samples were then immersed in 4% PFA and left on a rocker at 4 °C overnight. Following fixation, skin samples were trimmed and placed into cassettes and underwent the same washes as fixed lung tissue (detailed above). Skin samples were left in the final DEPC-treated 70% ethanol wash for tissue processing and embedding by the CCHMC Pathology Core. Samples were then sectioned for histology and immunofluorescence, conducted in the same manner as detailed above in the processing of fixed lung tissue.

### Trans-epidermal water loss (TEWL) measurements

TEWL was measured with the Dermalab instrument (cyberDERM; Cortex Technology, Media, Pa) as previously described^82^. The temperature and relative humidity of the room in which the measurements were taken varied between 22°C and 25.6°C and 36.4% and 48.1%, respectively. All TEWL measurements were taken after the mice had acclimated to the temperature and relative humidity of the room for at least 15 minutes. The probe was placed with very slight pressure on the center backside of the mouse following anesthetization with isoflurane. The rate of TEWL stabilized after 1min, and measurements were recorded as grams per meter squared per hour. TEWL was measured for the same cohort of mice at multiple time points: PND21, 28, 35, and 42. The same investigator measured TEWL on all mice at all timepoints.

### Influenza lung injury

Mice were administered PR8 H1N1 influenza at 8-10 weeks of age as previously described^81^. Multiple cohorts of mice were given either 16HAU, 20HAU, or 32HAU of PR8 H1N1 influenza via intranasal instillation. Following infection, mice were monitored daily for 28days, and lung injury was assessed via histology and immunofluorescence.

### Mouse intestinal harvest

Immediately following anesthetization via CO_2_ inhalation, the murine abdominal wall was opened, and the entire small intestine and colon were removed. The small intestine and colon were straightened to full length, and the small intestine was divided into fourths. The first section was collected as the duodenum, the third section as the jejunum, and the fourth section as the ileum (the second section was discarded). Each section was then divided again into fourths prior to collection, with the proximal-most subsection ultimately fixed in 10% formalin. Fecal contents were flushed from each subsection via blunt needle syringe, and each sample was subsequently flushed with 10% formalin. Each subsection of tissue was then cut longitudinally, laid flat onto a wax box filled with 10% formalin, and pinned by the edges. Intestinal samples collected at PND6 and 12 did not undergo flushing or longitudinal cutting due to size and fragility of the tissue. All samples were fixed overnight in 10% formalin at 4°C. The following day, samples were rinsed 3 times with 70% ETOH for 10min each.

### Agar blocking of intestinal tissue for histology

2% agar solution in DI H2O was placed in a 60°C water bath prior to blocking. 70% ETOH was removed from the wax box to allow the tissue samples to dry slightly. Each intestinal sample was fully covered with minimal 2% agar via plastic transfer pipette. After the agar cooled, it was first cut into rectangles around the tissue, then cut longitudinally to halve the tissue. Each tissue section was then placed such that the two halves faced inward. The paired halves of the duodenal, jejunal, and ileal samples were then placed back-to-back, covered in another layer of agar, and cooled. PND6 and 12 duodenal/jejunal/ileal samples were not halved in this manner and were immediately oriented back-to-back prior to the addition of the second agar layer. Excess agar was trimmed, and the agar block was placed into a cassette, which was then placed in a container with 70% ETOH. The CCHMC Pathology Core paraffin-embedded, sectioned, and provided H&E stains for these samples. PAS and Oil Red O staining obtained from Newcomer Supply (Middleton WI) and were performed per the manufacturer’s protocol.

### Intestinal morphometry measurements

Vacuole presence in intestinal villi was quantified based on the presence of any vacuoles in villi with intact tips and noted as low, medium, or severe grade. Crypt height was measured exclusively from well-oriented crypts from the base of the crypt to the base of the surrounding villi. Mean crypt height, villus height, and vacuole frequency were counted per 100 intact crypts or villi per sample if present in the histology; otherwise, all intact crypts or villi were counted per sample.

### Lung morphometry measurements

Morphometry methods to quantify alveolar structure were used as previously described^83^. The FIJI-macro provided in Salaets et al. was edited to allow jpg files for input. The proximal, intermediate, and distal fields of the left lobe of each lung sample were taken via brightfield imaging on the Nikon NiE upright microscope and saved as tiff files. Images were taken at 20X objective and at the same exposure. Prior to running the macro, image fields from all samples were batch-converted from tiff to jpg format. With the FIJI-macro, each image field was resized to 680×680 pixels, and non-parenchymal tissues (conductive airways, arteries, veins) were manually selected via the freehand selection tool. Smaller exudates were automatically deleted, and each image field was assessed for manual deletion of larger or connected exudates. The mask was automatically cleaned and smoothened, and intersections were counted in a grid comprised of 8 vertical and 8 horizontal lines. The number of intersections, alongside test points that fell on parenchymal (reference) tissue and test points that fell on alveolar septa, were provided as output for each image field. All lung morphometry parameters were calculated from these outputs.

## Supplemental Materials

**Supplemental Figure 1.**
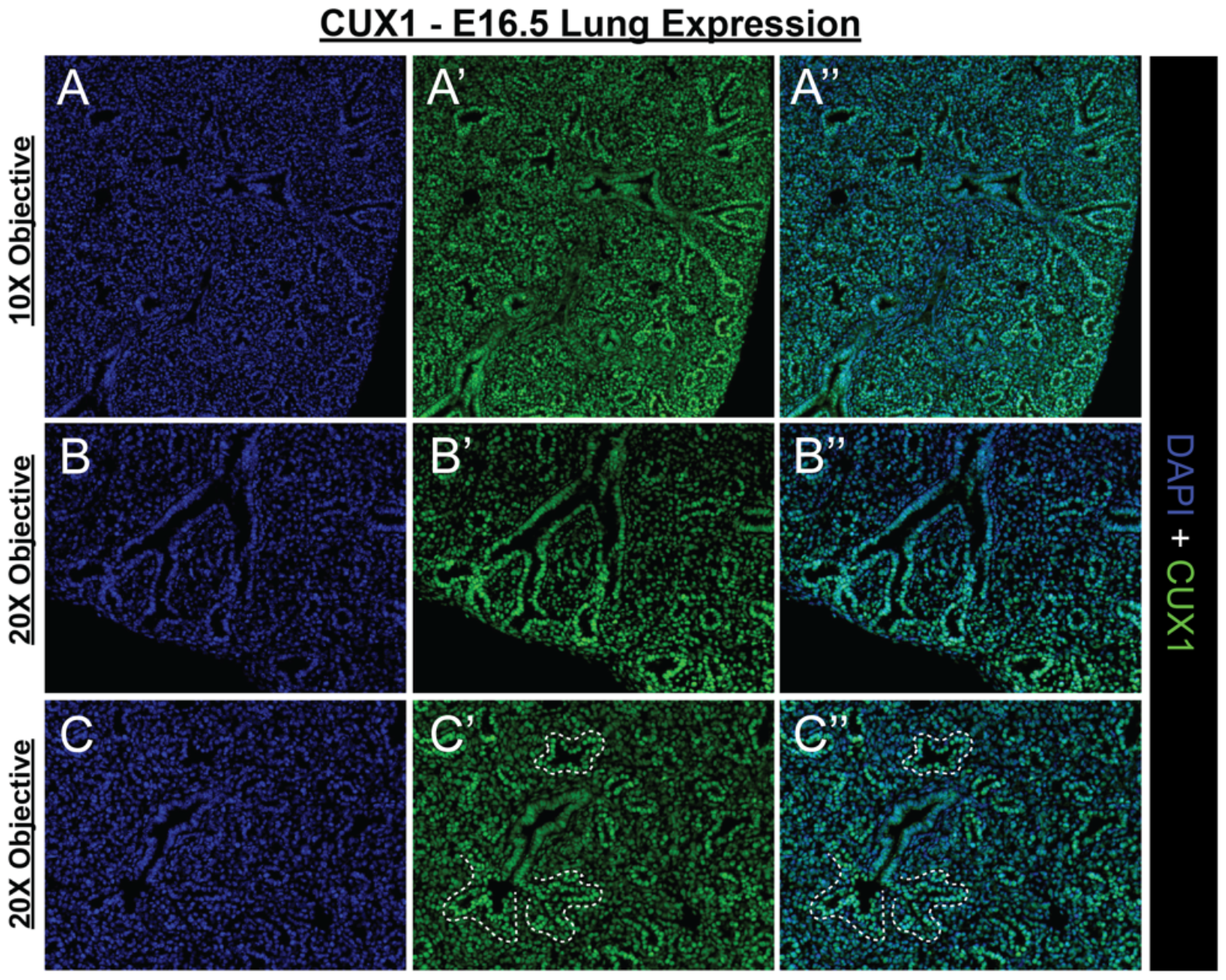
CUX1 protein is expressed in distal tips of the developing lung epithelium. (A-C) IHC of WT mouse lung demonstrating Cux1 protein expression at E16.5, with broad expression most pronounced in the distal lung epithelium.

**Supplemental Figure 2.**
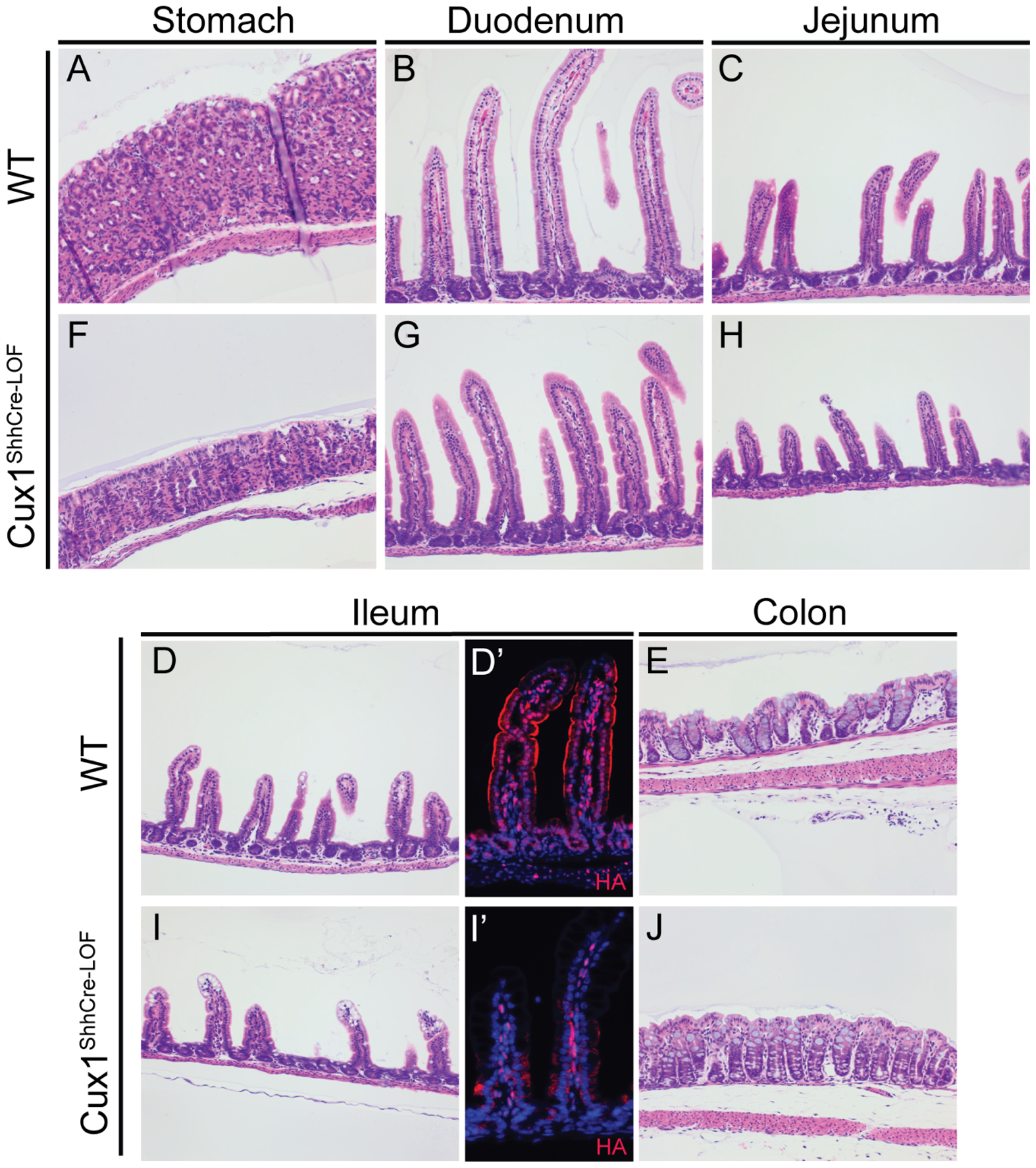
Evaluation of gut tube development in Cux1^ShhCre-LOF^ mice and WT littermates. No discernable differences were noted in the duodenum, jejunum, or colon of WT mice (A-E) compared to Cux1^ShhCre-LOF^ mice (F-J) at PND16. Vacuolization was noted in the ileum of Cux1^ShhCre-LOF^ mice (compare to Figure 7). Anti-HA immunostaining confirmed loss of epithelial CUX1-HA expression in Cux1^ShhCre-LOF^ ileum (D’ v I’).

**Supplemental Figure 3.**
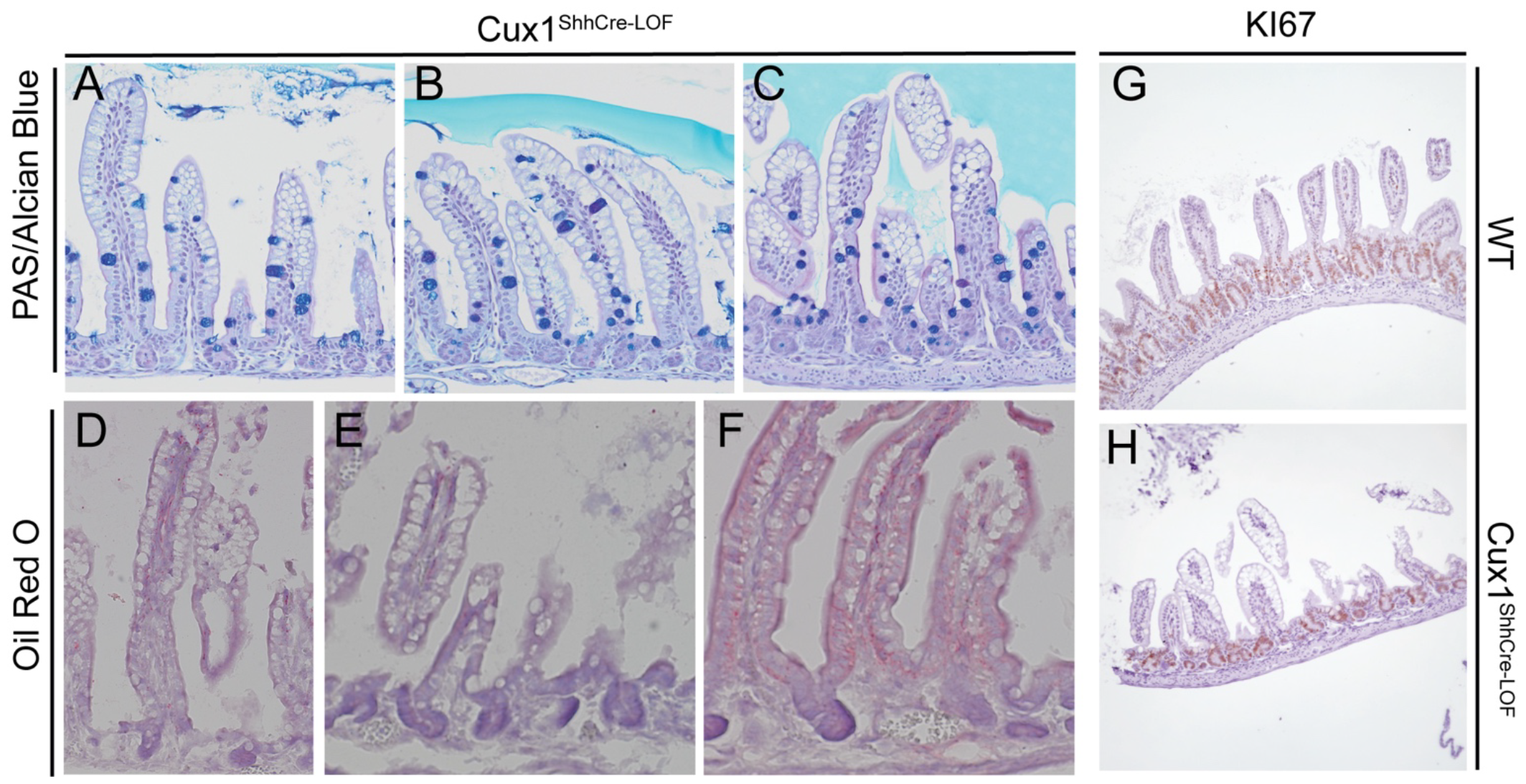
Ileal vacuoles present in Cux1^ShhCre-LOF^ mice do not contain mucus or lipids. (A-C) Alcian blue stain reveals mucous is not present in ileal vacuoles of Cux1^ShhCre-LOF^ mice. Goblet cells are stained blue and act as a positive control. (D-F) Oil Red O stain shows no lipids are present in the vacuoles of Cux1^ShhCre-LOF^ mice. (G-H) No difference in ileal crypt proliferation denoted by Ki67 expression in Cux1^ShhCre-LOF^ animals.

**Supplemental Figure 4.**
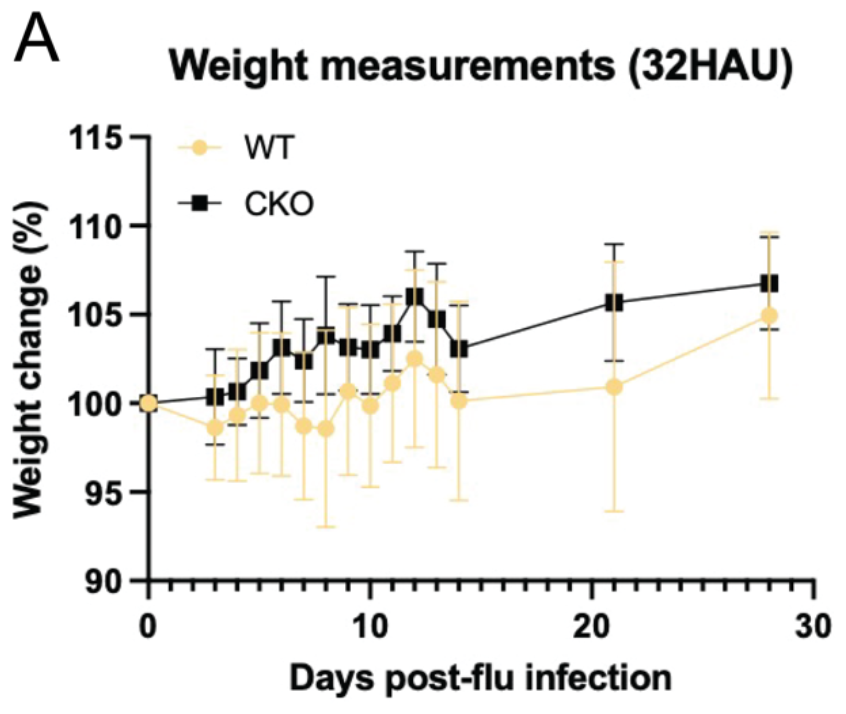
No difference in weight trend following influenza in Cux1^ShhCre-LOF^ mice.

**Supplemental Figure 5.**
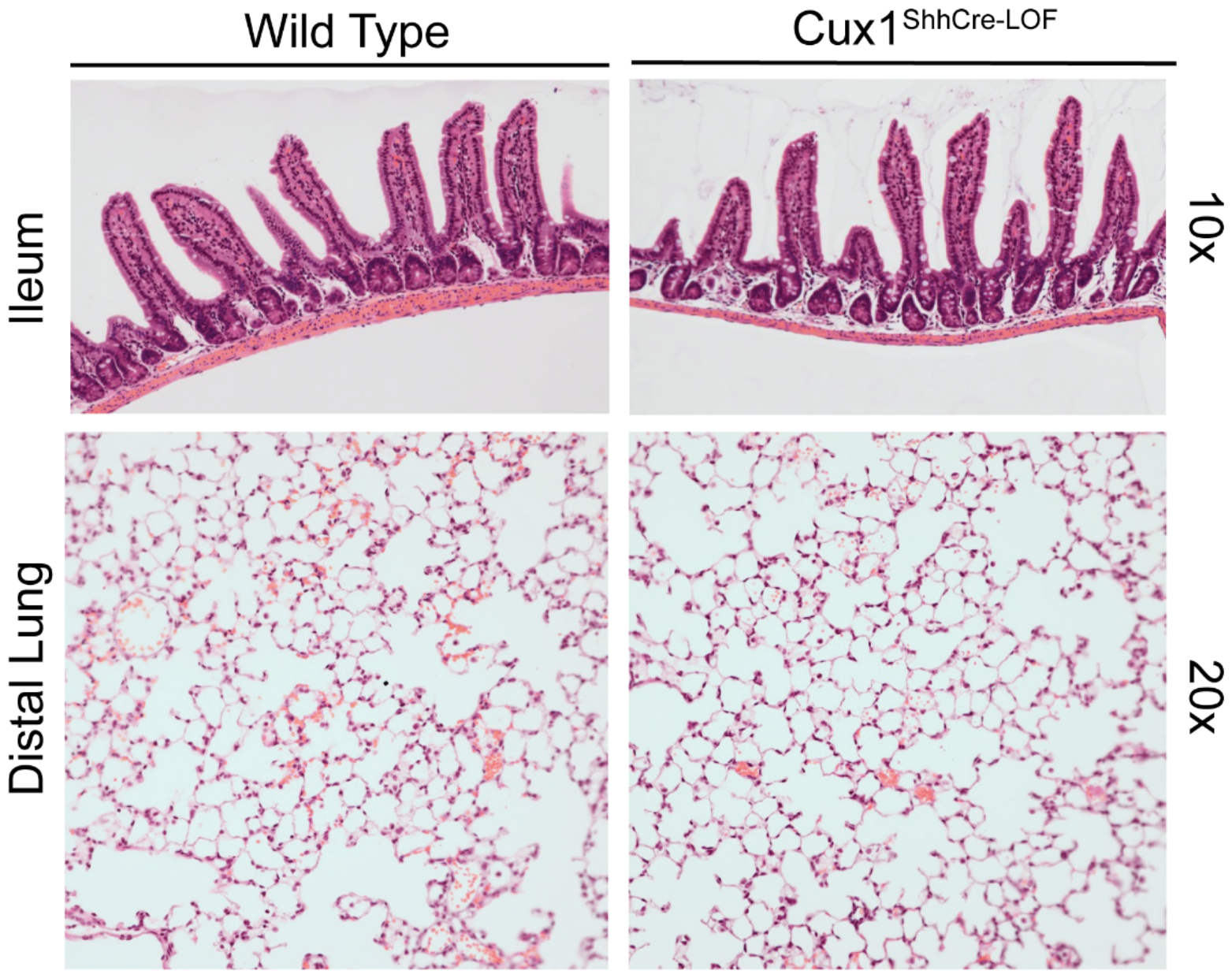
Lack of histological differences in either intestine or lung at outset of lung infection. At PND35, no discernable differences are notable in intestine or lung of Cux1^ShhCre-LOF^ mice.

